# APEX2 proximity labeling maps the mGBP2-proximal proteome at the *Toxoplasma gondii* parasitophorous vacuole

**DOI:** 10.64898/2026.10.13.758798

**Authors:** Imke Bradtmöller, Gereon Poschmann, Kai Stühler, Daniel Degrandi, Klaus Pfeffer

**Affiliations:** Institute of Medical Microbiology and Hospital Hygiene, Heinrich Heine University, Düsseldorf, Germany; Molecular Proteomics Laboratory, BMFZ, Heinrich-Heine-Universität, Düsseldorf, Germany

**Keywords:** APEX2, Proximity labeling, Apicomplexa, Parasite, Guanylate-binding proteins

## Abstract

Interferon-inducible guanylate binding proteins (GBPs) are large GTPases crucial for the host immune response against *Toxoplasma gondii*, a globally prevalent intracellular apicomplexan parasite. Upon infection, *T. gondii* forms a parasitophorous vacuole (PV) that protects the parasite from host intracellular defense mechanisms. Murine GBPs (mGBPs) can recognize these PVs and assemble into supramolecular homo- and heterocomplexes, which is an essential step for PV membrane disruption, ultimately enabling cell-autonomous immune control of vacuole-resident pathogens. Our previous work has shown that mGBP2 is an important factor in the immune control of *T. gondii*. To further elucidate the functions of mGBP2, we employed an APEX2-based proximity labeling strategy to biotinylate proteins in the vicinity of mGBP2 during *T. gondii* infection. Mass spectrometry analysis of the biotinylated proteins revealed several novel PV-associated proteins such as Cyclooxygenase 2 (COX2/ Ptgs2), Caspase 12 (Cas12), Annexin 11 (Anxa11), Calnexin (Canx), Galectin 3 (Gal3), and Transgelin (Tagln) and valosin-containing protein (VCP) that co-recruit with mGBP2 at *T. gondii* PVs. In addition, we investigated their regulation by IFN-γ and determined that inhibition of COX2 severely impairs *T. gondii* growth, highlighting a critical role for COX2 for parasite survival.

## INTRODUCTION

The innate ability of cells to resist invading pathogens is dependent on cell-autonomous immunity. Upon recognition of pathogens the production of Interferons (IFN) is a crucial step in the activation of cell autonomous immunity and necessary for the initial restriction of microbial replication (Man, Place et al. 2017, Kak, Raza et al. 2018). Among IFN-γ–induced Interferon-stimulated genes (ISGs) four major families of GTPases: the 45-kDa immunity-related GTPases (IRGs), VLIGs, myxovirus resistance (Mx) proteins and the 65- to 73-kDa guanylate-binding proteins (GBPs) (Boehm, Guethlein et al. 1998, Martens and Howard 2006, Degrandi, Konermann et al. 2007, Haller, Stertz et al. 2007, Zhao, Rohde et al. 2009, Hunn, Feng et al. 2011, Praefcke 2018) have been identified. Throughout the vertebrate linage GBPs show a high level of conservation (Kresse, Konermann et al. 2008). In humans seven hGBPs as well as at least one pseudogene located on chromosome 1 have been described (Man, Place et al. 2017, Ngo and Man 2017). The eleven Murine Gbp genes are located on chromosome 3 (*Gbp1*, *Gbp2*, *Gbp3, Gbp5* and *Gbp7*) and chromosome 5 *(Gbp4, Gbp6, Gbp8, Gbp9, Gbp10 and Gbp11)* respectively (Degrandi, Konermann et al. 2007, Kresse, Konermann et al. 2008, Man, Place et al. 2017).

GBPs exhibit a variety of distinct cellular functions, including antiparasitic, antiviral and antibacterial activity, as well as inhibiting cell proliferation and suppressing tumours (Degrandi, Konermann et al. 2007, Coers, Brown et al. 2018, Steffens, Beuter-Gunia et al. 2020)). Murine GBP2 has been shown to target intracellular pathogens like *Salmonella typhimurium* (Ngo and Man 2017) and exhibit antiviral activity against vesicular stomatitis virus and encephalomyocarditis virus (Carter, Gorbacheva et al. 2005). Furthermore, the deletion of genes Gbp1, Gbp2, Gbp3, Gbp5, and Gbp7 in mice is associated with a severe susceptibility to a broad range of pathogens including *Toxoplasma gondii* (*T. gondii*) (Yamamoto, Okuyama et al. 2012, Degrandi, Kravets et al. 2013, Selleck, Fentress et al. 2013, Steffens, Beuter-Gunia et al. 2020, Feng, Enosi Tuipulotu et al. 2022).

Several mGBPs such as mGBP1, mGBP2, and mGBP7 play an essential role for the resistance against *T. gondii*, an intracellular apicomplexan protozoan parasite by restricting the growth and mediating killing *T. gondii* in murine fibroblasts, macrophages and astrocytes in an IFN-γ dependant manner (Degrandi, Konermann et al. 2007, Yamamoto, Okuyama et al. 2012, Degrandi, Kravets et al. 2013, Kravets, Degrandi et al. 2016, Steffens, Beuter-Gunia et al. 2020).

Following the active invasion of host cells, *T. gondii* forms a parasitophorous vacuole (PV), a cytoplasmic membranous structure that envelops the invading *T. gondii* in the host cell cell (Matta, Rinkenberger et al. 2021).The PV serves as a replicative niche for the parasite (Sibley 2011). Upon induction of IFN-γ, mGBP2 has been observed to assemble into vesicle-like structures (VLS) within the cytoplasm. Through an as-yet-unknown IFN-γ–dependent mechanism, murine GBPs (mGBPs) can specifically recruit to and accumulate on pathogen-containing vacuoles, thereby promoting cell-autonomous clearance of the pathogen (Degrandi, Kravets et al. 2013, Lindenberg, Molleken et al. 2017, Steffens, Beuter-Gunia et al. 2020). Following induction of IFN-γ, mGBP2 has been observed to assemble into vesicle-like structures (VLS) within the cytoplasm. After invasion of *T. gondii* into the host cells, mGBP2 directly attacks the PV membrane (PVM) by the formation of supramolecular complexes leading to the disruption of the PVM (Degrandi, Kravets et al. 2013, Kravets, Degrandi et al. 2016). This process is followed by further recruitment of mGBP2 to the plasma membrane of the intracellular parasite leading to its destruction (Kravets, Degrandi et al. 2016). Using multiparameter fluorescence image spectroscopy (MFIS) it has been shown that mGBP2 can form homo- or heteromultimers with mGBP1, mGBP3 and mGBP5 within VLS and at the PVM (Kravets, Degrandi et al. 2016) Moreover, it has been demonstrated that both mGBP2 and mGBP7 are found at the parasite plasma membrane within the targeted parasite (Kravets, Degrandi et al. 2016, Steffens, Beuter-Gunia et al. 2020). Recently, we identified Galectin-9 (Gal9) as a critical mGBP2 interaction partner that is engaged by mGBP2 for immunity against *T. gondii*. To continue the search for mGBP2 associated proteins which are important for cell autonomous immunity, in this study, a proximity labeling approach was performed.

In recent years, proximity labeling strategies have shown their utility in studying protein-protein interactions (PPIs) and molecular compartments (Qin, Cho et al. 2021). Proximity labeling approaches offer several distinct advantages over previous strategies for elucidating the molecular environment of mGBPs. On the one hand, proximity labeling can be performed in living cells, which enables the identification of endogenous PPIs. On the other hand, these strategies are designed to specifically label the proximity proteome of target proteins, including proteins that do not interact with the target protein directly (Qin, Cho et al. 2021). Ascorbate peroxidase 2 (APEX2) is a proximity labeling catalyst that enables efficient biotinylation in only one minute making it particularly advantageous for detecting also weak and transient interactions (Lam, Martell et al. 2015).

The aim of this study was to further explore the molecular environment of mGBP2 at the *T*. *gondii* PV. Using an APEX2-based proximity labeling approach we identified by mass spectrometry (MS) analysis a novel set of biotinylated proteins in proximity to mGBP2 in IFN-γ stimulated and *T. gondii* infected cells. Interestingly, these proteins, including COX2, co-recruit with mGBP2 to the PV. Furthermore, we investigated the influence of IFN-γ stimulation on these novel proteins and demonstrated that inhibition of one of those, COX2, in contrast to mGBP2, leads to impaired *T. gondii* growth indicating a crucial role of COX2 for parasite survival.

## MATERIALS AND METHODS

### Expression construct

The WT ORF of mGBP2 (NCBI accession numbers: mGBP2, NM_010260.1) was synthesized into the pWPXL plasmid as N-terminal APEX2-fusion construct (Genscript, Leiden, Netherlands). pWPXL was a gift from Didier Trono (Addgene plasmid # 12257; http://n2t.net/addgene:12257; RRID:Addgene 12257). The lentiviral envelope vector pLP/VSVG (Invitrogen) and the packaging vector psPAX2 (Trono lab) were used for the lentiviral genetic transfer.

### Cell culture and transduction

WT (C57BL/6N) and mGBP2^−/-^ murine embryonic fibroblasts (MEFs), human foreskin fibroblasts (Hs27, ATCC CRL-1634) and 293FT cells (Gibco, ThermoFisher; R700-07, USA) were cultured in Dulbecco’s modified Eagle’s medium (DMEM, Invitrogen/Gibco, USA) supplemented with 10% (v/v) heat-inactivated low endotoxin fetal bovine serum (FBS, Cambrex, USA), and 0.05 mM β-mercaptoethanol (Invitrogen/Gibco, USA). All recombinant lentiviruses were produced by transient transfection of 293FT cells according to the jetPRIME^®^ transfection protocol (Polyplus, USA). MEFs were seeded in 24 well plates (Corning Incorporated, Germany) and transduced with 500 µL of lentivirus with 25 µg Polybrene (Merck Millipore, Germany). After 6 h of incubation medium was changed. The transduction efficacy was analyzed by flow cytometry. Subsequently, only mCherrry or double GFP and mCherry positive cells were sorted and cultivated.

### In vitro passage of *T. gondii*

Tachyzoites from *T*. *gondii* strain ME49 (ATCC 50611) and *T. gondii* ME-GFP-Luc (Boyle, Saeij et al. 2007) were maintained by serial passage in confluent monolayers of HS27 cells.

### Infection of MEFs with *T*. *gondii*

Cells were stimulated with 200 U/mL IFN-γ (R&D Systems, Minneapolis, MN) 16 h before infection. For immunofluorescence, MEFs were cultured in 24-well plates (Falcon, BD Biosciences, Germany) on cover slips (ø 13 mm, VWR International, Germany) and infected with freshly harvested *T*. *gondii* with a MOI of 40:1.

### RNA isolation

1×10^5^ MEFs were seeded in 6 well plates. Cells were left untreated or treated with 200 U/mL IFN-γ (R&D Systems, Minneapolis, MN) 16 h or 500 ng/ml LPS (*Escherichia coli* O127:B8, Sigma-Aldrich, Merck KGaA, Germany) for 16h. Cells were then left uninfected or infected with freshly harvested *T*. *gondii* with a MOI of 40:1 for 2.5 hours. RNA isolation was performed using the RNeasy Plus Mini Kit (Qiagen, Germany) according to the manufacture’s instructions. RNA was eluted in 40 µL UltraPure™ DNAse/RNAse-Free distilled Water and concentration was measured using a NanoDrop device (ThermoFisher Scientific, USA).

### cDNA synthesis

cDNA synthesis was performed by mixing 1 µg of isolated RNA with 0.5 mM dNTPs, 0.5 µg Oligo(dT) (Invitrogen, ThermoFisher Scientific, USA), 1 µL RNase OUT (Invitrogen, ThermoFisher Scientific, USA) and up to 14 µL UltraPure™ DNAse/RNAse-Free Distilled Water (Invitrogen, ThermoFisher Scientific, USA). The mix was incubated at 65°C for 5 min. Then 1X First-Strand Buffer, 100 mM DTT, and 200 U SuperScriptTM III Reverse Transkriptase (Invitrogen, ThermoFisher Scientific, USA) was added. cDNA synthesis was performed in a Biometra Thermocycler T-Gradient ThermoBlock (Biometra, Germany) by incubation at 50°C for 90 min and 70°C for 15 min.

### Single clone generation

To generate single clone cell lines carrying the transduced Apex2-mGBP2 plasmid, the transduced cells were treated with trypsin for cell separation and counted using a Neubauer counting chamber. In a 96 well plate (Invitrogen, ThermoFisher Scientific, USA) cells were seeded with a calculated amount of 0.3 cells per well. After sufficient growth, the cells were expanded in cell culture flasks (Corning, USA) and the Apex2-mGBP2 expression was confirmed via PCR after RNA isolation and cDNA synthesis.

### Polymerase chain reaction (PCR)

To confirm the expression of Apex2-mGBP2, a PCR using the following primers was performed: Apex2-mGBP2fwd (GGCTGGCGTTGTTGCCGTTG) and Apex2-mGBP2rev (AGCGGAATCGTCTACCCCACT) was performed. The reaction mix contained 1x Green Go Taq Reaction Buffer (Promega, Germany), 10 mM dNTPs, 10 µM forward and reverse primer respectively, 1 µg of cDNA and 5 U/µL GoTaq DNA polymerase (Promega, USA). The reaction was performed in a Biometra Thermocycler T-Gradient ThermoBlock (Biometra, Germany) using the following cycles: denaturation 94°C for 30 s, then 30 cycles denaturation at 94 °C for 30 s, annealing at 58°C for 30 s and extension at 68 °C for 60 seconds, followed by a completion step at 68 °C for 300 s.

### Immunofluorescence analysis

Cells were fixed in 4% paraformaldehyde (PFA) (Sigma-Aldrich, Merck KGaA, Germany) permeabilized with 300 µL 0.02% saponin (Sigma-Aldrich Solutions, Merck KGaA, Germany) in PBS (Gibco, ThermoFisher, USA) at room temperature and blocked in 300 µL 0.002% saponin with 2% goat serum (DaKoCytomation, Denmark) in PBS at room temperature. The cell membrane of *T. gondii* was visualized by an anti-SAG1 antibody (Invitrogen #MA5-18268, ThermoFisher Scientific, USA) at a dilution of 1:200 in 0.0002% saponin with 2% goat serum in PBS. Afterwards three washing steps using 500 µL wash buffer (0.0002% saponin in PBS) was performed. As secondary reagents, a dilution of 1:200 Alexa Fluor^TM^ 633-conjugated goat anti-mouse IgG (H+L) (Invitrogen # A21050, ThermoFisher Scientific, USA) in 0.0002% saponin with 2% goatserum in PBS was used. Again, three washing steps using 500 µL wash buffer were performed. The nuclei were counterstained with a dilution of 1:2000 4’,6-diamidino-2-phenylindole (DAPI, Invitrogen, ThermoFisher Scientific, USA) in PBS. Staining of biotinylated proteins was performed using a 1:2000 dilution of Alexa-Fluor-568 dye-coupled streptavidin, which can bind biotinylated proteins, as an antibody substitute, in PBS. After another washing step using 500 µL wash buffer, the cover slips were fixed in fluorescence mounting medium (Fluoromount-G, Southern Biotechnology Associates, Birmingham, AL). Fluorescence was visualized using a LSM780 confocal microscope (Zeiss, Germany). Image analysis and processing was performed with the ZENblue 2.5 (Zeiss, Switzerland) software.

### APEX2 proximity labeling

To label nearby proteins with APEX2, 1×10^4^ MEF cells expressing the APEX2-mGBP2 fusion protein were seeded the day before in a 24-well plate (Merck KGaA, Germany). For fixation with subsequent staining, sterile cover slips were used analogous to immunofluorescence analysis. For subsequent following Streptavidin pulldown 5×10^4^ MEF cells expressing the APEX2-mGBP2 fusion protein were seeded in a 25 cm² cell culture flask instead. For examination during *T. gondii* infection, cells were stimulated 16 h beforehand with 200 U/ml IFN-γ and infected the next day with *T. gondii* at a cell-to-parasite ratio of 1:50 for 2.5 h. Biotin-Phenol (BP) (Sigma-Aldrich Solutions, Merck KGaA, Germany) was then added up to a final concentration of 1 mM to the medium and incubated at 37°C for 30 min. Subsequently, hydrogen peroxide (H_2_O_2_) (Sigma-Aldrich Solutions, Merck KGaA, Germany) was added up to a final concentration of 1 mM to initiate the reaction leading to a reactive biotin-phenoxyl species. After one minute, the medium was removed and the reaction was immediately stopped with 300 µL for immunoflourescence or 5 ml for Streptavidin pulldown quencher solution (10 mM sodium azide (Sigma-Aldrich Solutions, Merck KGaA, Germany), 10 mM sodium ascorbate (Sigma-Aldrich Solutions, Merck KGaA, Germany), and 5 mM Trolox (Sigma-Aldrich Solutions, Merck KGaA, Germany) in PBS. Subsequently, the cells were washed four more times with 300 µL or 5 ml quencher solution and three times with 300 µL or 5 ml PBS.

### Preparation of total cell lysates from eukaryotic cells

To prepare whole-cell extracts, a range of 1×10^6^ to 1×10^7^ cells were subjected to trypsinization, followed by washing with PBS. The cells were then resuspended in 100 µL cell lysis buffer (10 mM MgCl_2_ (Sigma-Aldrich Solutions, Merck KGaA, Germany), 10 mM KCl (Sigma-Aldrich Solutions, Merck KGaA, Germany), 50 mM HEPES (Serva Electrophoresis Gmbh, Germany), 10 % (v/v) Gylcerin (EMSURE®, Merck KGaA, Germany), 200 mM NaCl (Sigma-Aldrich Solutions, Merck KGaA, Germany), 0,5 % (w/v) NP-40 (Sigma-Aldrich Solutions, Merck KGaA, Germany), 1 mM DTT (Sigma-Aldrich Solutions, Merck KGaA, Germany) per 1 × 10^6^ cells supplemented with 1 mM quencher solution and incubated on a rotating wheel at 4°C for 20 minutes. After centrifugation at 13.000 rpm and 4 °C for 20 minutes, the supernatant was transferred to a fresh reaction tube, while the remaining cellular debris was discarded.

### Streptavidin pulldown

For the pulldown, 40 µL of the lysate was supplemented with 10 µL loading buffer (5x) (Boster Biological Technology, US), incubated at 95 °C for 5 minutes and used as control. 30 µL streptavidin sepharose beads (Roche, Switzerland) were washed with 300 µL cell lysis buffer and incubated with the remaining lysate (560 – 660 µL) over night at 4 °C with gentle rotation. After centrifugation at 3000 rpm for 2 minutes 40 µL of the supernatant were supplemented with 10 µL loading buffer (5x) and incubated at 95 °C for 5 minutes as flowthrough control. The beads were then washed twice with 300 µL 0.2% SDS (Serva Electrophoresis Gmbh, Germany) followed by washing steps once with 300 µL buffer 1 (20 mM Tris (Carl Roth GmbH + Co. KG, Germany) and 2% SDS, pH7.5), twice with 300 µL buffer 2 (0.1% DOC (Sigma-Aldrich Solutions, Merck KGaA, Germany), 1% Triton X-100 (Sigma-Aldrich Solutions, Merck KGaA, Germany), 500 mM NaCl, 1mM EDTA (Sigma-Aldrich Solutions, Merck KGaA, Germany) and 50 mM HEPES, pH7.5), once with 300 µL buffer 3 (250 mM LiCl (Sigma-Aldrich Solutions, Merck KGaA, Germany), 0.5% NP-40, 0.5% DOC, 1 mM EDTA and 10 mM Tris, pH8.1), twice with 300 µL buffer 4 (50 mM Tris and 50 mM NaCl, pH7.4) and finally twice with 300 µL PBS. The beads were then sedimented by centrifuging at 3000 rpm for 2 minutes, the supernatants removed and the beads subsequently resuspended in 40 µL SDS loading buffer (Boster Biological Technology, US) supplemented with 1 mM biotin (Sigma-Aldrich Solutions, Merck KGaA, Germany). They were carefully vortexed and spun down shortly followed by an incubation at 95 °C for 5 minutes. The beads were then centrifuged at 13,000 rpm for 1 minute and the supernatant was transferred into a fresh 1.5 ml tube.

### Sodium dodecyl sulfate–polyacrylamide gel electrophoresis

After streptavidin pulldown, the samples were incubated with loading buffer at 95°C for SDS Page as previously described. 25 µL of the protein sample was loaded on a 4-12% Bis-Tris-Gel (Invitrogen, Thermo Fisher Scientific, USA) which was placed in a XCELL SureLock Mini-Well gel chamber (Invitrogen, Thermo Fisher Scientific, USA) filled with 1x NuPAGE^TM^ MES SDS Running buffer (Invitrogen, Thermo Fisher Scientific, USA). Five µL of Precision Plus Protein Dual Color Standards (Bio-Rad Laboratories, #1610374, Germany) was used as molecular weight ruler. The gel was run at 120-150 V for 1 h followed by western blot analysis.

### Western Blot

The proteins were transferred to a 0.2 µm Nitrocellulose membrane using the Bio-Rad Trans-Blot Turbo Transfer System using the Transfer Pack mini gel 7×8,5 (Bio-Rad Laboratories, Germany) following the manufacturers manual. The 1.5 MM Gel protocol was used running at

1.3 A constant for 10 min. The membrane was then blocked by incubation in 5% skim milk powder (Merck Millipore, Germany) or 5% BSA (Sigma-Aldrich, Merck KGaA, Germany) depending on used antibody for 1 hour. Primary antibody (rabbit anti-mCherry antibody (E5D8F) (Cell Signaling #43590) (1:2500), mouse anti-COX2 antibody Invitrogen #35-8200) (1:500) was diluted in 5 % MP and incubated overnight at 5 °C. Secondary antibody (anti-rabbit HRP-linked antibody (Cell Signaling #7074) (1:1000), anti-mouse HRP-linked antibody (Cell Signaling #7076) (1:1000) was diluted in 3 % MP and incubated for 1 hour. Streptavidin-HRP (Thermo-Fisher #SA10001) (1:1000) was diluted in 5% BSA and incubated overnight at 5 °C. Chemiluminescence detection was performed using SuperSignal™ West Pico PLUS Chemiluminescence substrate (Thermo Fisher Scientific, USA) at the ECL Imager Chemostar Intas (Science Imaging Instruments, Germany).

### Liquid chromatography coupled mass spectrometric analysis of proteins enriched after proximal labeling

Eluates from streptavidin mediated enrichment of biotinylated proteins from murine embryonal fibroblasts (4 groups with each n=5 independent replicates: without biotin phenol and without *T. gondii* infection, without biotin phenol and with *T. gondii* infection, with biotin phenol and without *T. gondii* infection, with biotin phenol and with *T. gondii* infection) were prepared for mass spectrometric analysis by in-gel digestion as described (Poschmann, Prescher et al. 2021). Briefly, samples were briefly stacked into a polyacrylamide gel, the gel was stained with Coomassie brilliant blue and protein containing bands cut out, proteins reduced and alkylated with iodoacetamide and finally digested with trypsin. Resulting peptides were extracted from the gel and one third of the sample analyzed by liquid chromatography coupled mass spectrometry.

First peptides were separated by a one-hour gradient on 25 cm long C18 columns on an Ultimate3000 rapid separation liquid chromatography system (Thermo Fisher Scientific) essentially as described previously (Poschmann, Schwermer et al. 2026). Then, mass spectrometric analysis was carried out on an online - via nano-electrospray coupled - Orbitrap Fusion Lumos instrument (Thermo Fisher Scientific) operated in data-dependent, positive mode. For each acquisition cycle, one survey spectrum was recorded in the orbitrap analyser (resolution 120000, scan range 350 – 2000 m/z, maximum injection time 50 ms, AGC target 400000, profile mode). Subsequently, 2-6fold charged precursors were selected in the quadrupole (isolation window 1.6 m/z), fragmented by higher energy collisional dissociation (35% collision energy) and fragment spectra recorded in the ion-trap (scan rate rapid, scan range auto, maximum injection time 35 ms, AGC target 10000, centroid mode). Cycle time was one second and dynamic exclusion set to one minute.

The resulting protein table was further analyzed within the Perseus 1.6.6.0 framework (Max Planck Institute for Biochemistry, Planegg, Germany). Potential contaminants, reverse hits, proteins identified “by site” and proteins identified with only one peptide were removed from the data matrix and only proteins considered with at least four valid values in the sample group of samples to which biotin phenol was added. Normalized intensities were log_2_ transformed to reach a normal-like distribution of quantitative values. Missing values were imputed with random values drawn from a down-shifted normal distribution (0.3 SD width and 1.8 SD down-shift) to account for the high number of missing values in the group of samples which has not be treated with biotin phenol to initiate APEX2 mediated labeling. The significance analysis of microarrays method (Tusher, Tibshirani et al. 2001), based on Student’s t-tests and a permutation-based control of the false discovery rate (5%, S_0_ was 0.6), was used for the comparison of pairs of sample groups with a minimum of initially four valid values in at least one of samples groups in the respective comparison.

The mass spectrometry proteomics data have been deposited to the ProteomeXchange Consortium via the PRIDE (Perez-Riverol, Bandla et al. 2025) partner repository with the dataset identifier PXD076822.

### *T. gondii* proliferation assay

To evaluate the effect of COX inhibitors on *T. gondii* proliferation, in vitro inhibition assays were conducted. 4×10^4^ Hs27 or MEF cells were seeded in 96-well microtiter plates (Invitrogen, ThermoFisher Scientific, USA) with a total culture volume of 200 µL each well. To avoid edge effects, only the inner 60 wells of each plate were used. The host cells were then infected with 1×10^4^ Luciferase expressing tachyzoites per well, with a MOI ratio of 0.25:1. Simultaneously the inhibitors were added to the cultures at various concentrations (ranging from 200 µM to 1.25 µM). As controls, *T. gondii* infected cells only and human IFN-γ (300 U/ml) (Merck, Darmstadt, Germany) or murine IFN-γ (200 U/mL) (R&D Systems, Minneapolis, MN) pre-stimulated and infected cells were used. To quantify the proliferation of *T. gondii* the medium was removed after 48 h and the cells were washed with PBS before 40 µL 1x passive lysis buffer (Promega, USA) was added and incubated for 1 h. 20 µL of the respective lysate was transferred to a white flat-bottom 96-well plate (Sigma-Aldrich, Merck KGaA, Germany). Luciferase activity was measured in a Berthold Mithras LB 940 multimode plate reader (Berthold Technologies, Germany) by injecting 50 µL Luciferase substrate (Promega, USA) and using 2 s integration time. The IC_50_ values (minimal concentration of compounds required for 50% inhibition *in vitro*) were determined by non-linear regression analysis through the statistics software GraphPad PRISM^TM^ (Version 9.3.1; San Diego, USA).

### MTT Assay

The cytotoxic effect on Hs27 host cell viability exhibited by the COX inhibitors was measured by performing MTT (3-(4,5-di methyl thiazol-2-yl)-2,5-diphenyltetrazolium bromide) assays using the CyQUANT™ MTT Cell Viability Assay (Invitrogen, Thermo Fisher Scientific, USA). 4×10^4^ cells per well were seeded in a 96 well plate (Invitrogen, ThermoFisher Scientific, USA) and the inhibitor solved in DMSO was added in serial dilution starting at 200 µM 48 h hours before start of the assay. Serial dilution of DMSO was used as control. Read out was performed in a Tecan Sunrise microplate reader at 570 nm using the Magellan software (Tecan, Switzerland). The half maximal cytotoxic concentration (CC_50_ values) of each inhibitor against Hs27 host cells relative to DMSO treated samples were determined using GraphPad PRISM^TM^ statistics software (Version 9.3.1; San Diego, CA).

### q-PCR

To investigate the fold change of selected target transcripts after IFN-γ/LPS stimulation and/or *T. gondii* infection, RT PCR was performed. The reaction mix was prepared by combining 12.5 µL of Takyon® No ROX SYBR 2X MasterMix blue dTTP (Kaneka Eurogentec S.A., Belgium), 2.5 µL of UltraPure™ DNA/RNAse-Free Distilled Water, and 1 µM of each respective forward and reverse primer. 20 µL of the reaction mix was added into each well in a qPCR plate (Azenta, USA) and 25 ng of the cDNA was added to a total volume of 25 µL. RT-PCR was performed in a Biometra Thermocycler T-Gradient ThermoBlock (Biometra, Germany) using the following settings: 95°C for 15 min, then 40 cycles of 95°C for 15 sec followed by 60°C for 1 min, a final melt curve from 65°C to 95°C in 0.5°C steps each for 15 sec was carried out at the end. β-actin served as an internal standard to normalize the amount of cDNA across samples. The difference in cycle threshold (ct) values (Δct) between the ct-value of the target gene and the ct-value of β-actin was calculated for each sample. To calculate the relative fold change, the Δct value of the sample was subtracted from the respective Δct value of the unstimulated control group (ΔΔct), followed by the calculation: fold change = 2^−ΔΔct^.

## RESULTS

### APEX2-mGBP2 is able to recruit to *T. gondii* PVs after stimulation and biotinylates proteins in proximity

To identify the molecular environment of mGBP2 we designed a construct for an Apex2-mGBP2 fusion protein expressed under the EF1α promoter. Via lentiviral transduction this construct was introduced into mGBP2^−/-^ MEFs cells and single-cell cloning of the transduced cells was performed via limiting dilution. After sufficient growth we isolated RNA from the single clones, performed cDNA synthesis and then performed PCR for APEX2-mGBP2 expression using primers specific for APEX2 and mGBP2.

Next, we verified the correct subcellular localization of the introduced fusion protein. Previous studies have shown that mGBPs, including mGBP2, recruit to the PV of *T. gondii* upon IFN-γ stimulation (Degrandi, Konermann et al. 2007, Kravets, Degrandi et al. 2012, Kravets, Degrandi et al. 2016).To investigate whether APEX2-mGBP2 retains PV recruitment and whether the APEX2-catalyzed labeling reaction occurs as expected, we performed APEX2 proximity labeling in unstimulated and IFN-γ stimulated as well as *T. gondii* infected and uninfected cells expressing APEX2-mGBP2. To visualize the labeled proteins, we used Streptavidin fused to Alexa568. In the uninfected cells, Streptavidin binding was seen in the cytosol where GBPs usually reside in vesicle-like structures (Degrandi, Konermann et al. 2007) (Fig. 1A). Untreated cells showed background staining of presumably endogenously biotinylated proteins which might be attributed to proteins like the *T. gondii* Acetyl-coA-carboxylase and pyruvate carboxylase which have been shown to be located in the apicoplast and mitochondria respectively (Blackman and Bannister 2001, Jelenska, Crawford et al. 2001) (Fig. 1B). Upon IFN-γ stimulation a distinct accumulation of biotinylated proteins was detected at the *T. gondii* PV, indicating correct localization of the APEX2-mGBP2 protein and successful biotin labeling of proximal proteins (Fig. 1C/D).

**Figure 1.**
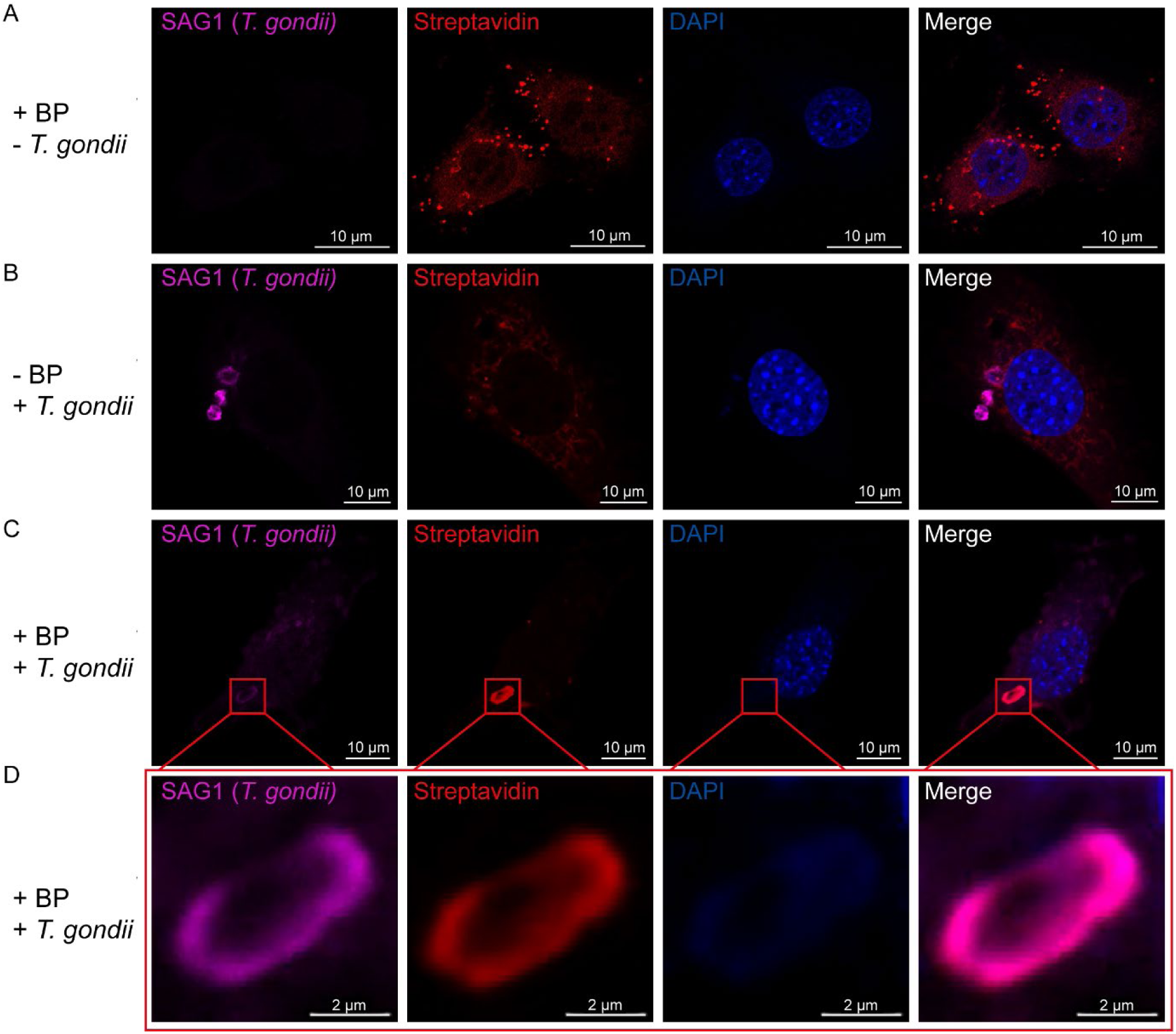
Validation of APEX2-mGBP2 mediated proximity labeling (PL) by confocal microscopy. mGBP2-/- MEFs expressing an APEX2-mGBP2 fusion protein were used for PL after *T. gondii* infection. Cells were treated with 200 U IFN-γ 16 h before infection with *T. gondii* ME49 or left untreated / uninfected as indicated. Biotinylated proteins were stained with Streptavidin AF568, T*. gondii* with anti-SAG1/anti-mouse AF633. Nuclei were stained with DAPI. Slides were analyzed by confocal microscopy. Bars, 10 μm (A-C), 2 µm (D).

### Proximity labeling reveals the molecular environment of mGBP2 at the *T. gondii* PV

Following these experiments, we repeated APEX2-proximity labeling in IFN-γ stimulated cells that were either uninfected or infected with *T. gondii* and treated with BP or left untreated. We then enriched the biotinylated proteins for further analysis via streptavidin beads. Western blot showed only minor signals of biotinylated proteins in the sample without biotin phenol incubation while the treated samples show a broad span of APEX2 biotinylated proteins of various molecular weights (Fig. 2). Enriched proteins of five independent replicates per group were then collectively analyzed by quantitative mass spectrometry.

**Figure 2.**
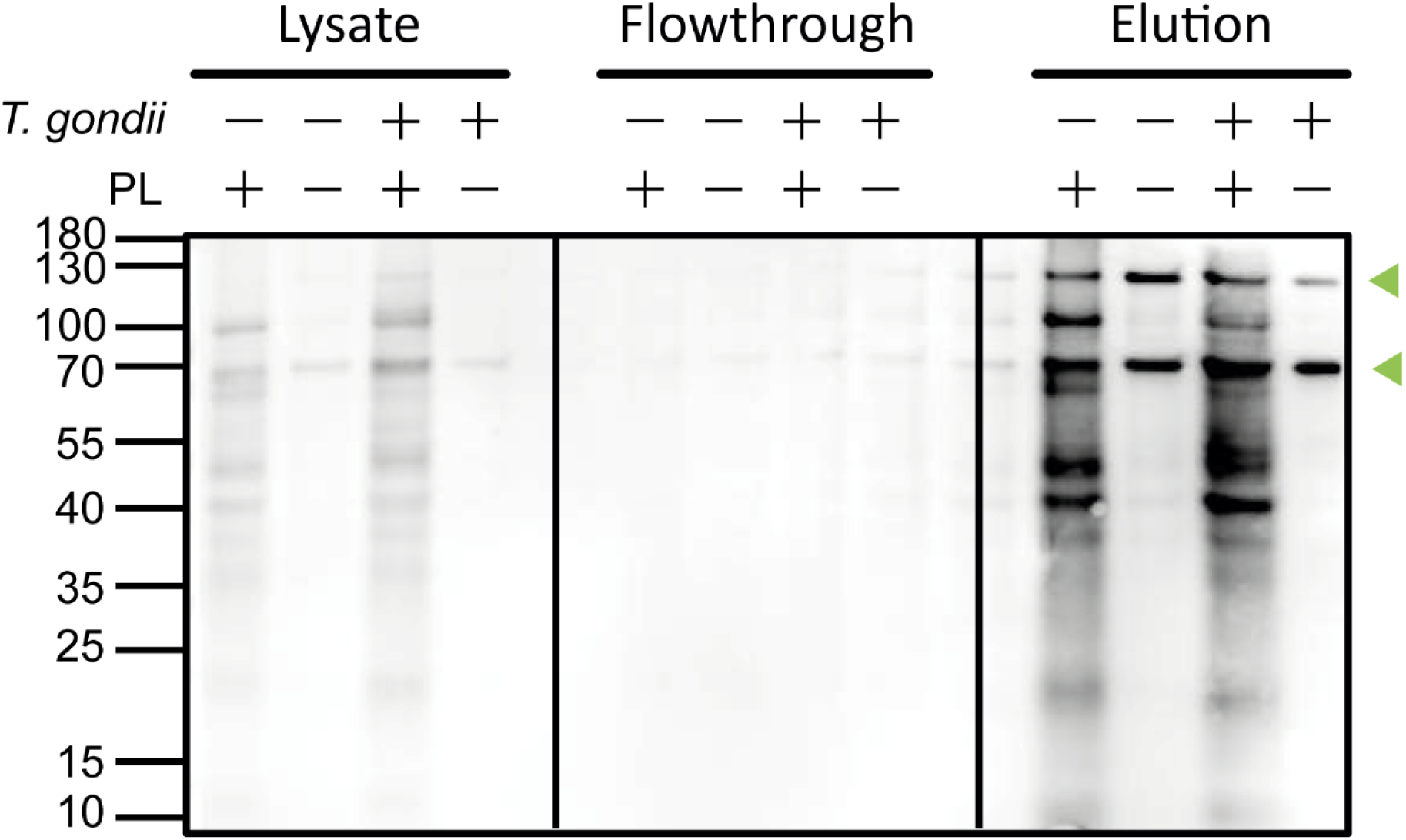
Streptavidin-HRP Western blot analysis of APEX2 proximity labeling in *T. gondii* infected MEFs. Cells were treated with 200 U/ml IFN-γ and left uninfected or were infected with *T. gondii* ME49 (*T. gondii*), incubated with biotin phenol or left untreated. Molecular mass standards are shown in kDa on the left side of the figure. Arrows indicate presumably endogenously biotinylated proteins

Here, we found a strong increase of protein abundance after induction of biotinylation by biotin phenol as well as a distinct clustering of protein abundance data depending on T. gondii infection (Suppl. Fig. S2A). Among the 294 higher abundant proteins in biotin phenol treated samples after *T. gondii* infection, we discovered besides mGBP2 also other mGBPs including mGBP6 and mGBP5 but also known interacting GBPs, such as mGBP1 and mGBP3 (Kravets, Degrandi et al. 2016). Furthermore, we found proteins that are already known to directly interact with mGBP2 including Irgb6, which plays a crucial role in mGBP2 loading of the *T. gondii* PV (Lee, Yamada et al. 2020). Interestingly, we also identified proteins that are known to recruit to the *T. gondii* PV such as Irga6 (Lee, Yamada et al. 2020), p62 (Lee, Sasai et al. 2015), PDCD6IP/ALIX (Rivera-Cuevas, Mayoral et al. 2021), RNF213 (Hernandez, Walsh et al. 2022), indicating success of the proximity labeling experiment and validating our results.

We decided to further investigate proteins that specifically were labeled in T. gondii infection in high amounts but have not been described in that context yet, which included the Cyclooxygenase 2 (COX2/ Ptgs2), Caspase 12 (Cas12), Annexin 11 (Anxa11), Calnexin (Canx), Galectin 3 (Gal3), Transgelin (Tagln), and valosin-containing protein (VCP), a member of the adenosine triphosphatase family (Suppl. Fig. S2B)

### APEX2-labeled proteins recruit to *T. gondii* PVs together with mGBP2

For each protein of interest, we generated constructs expressing the gene of interest fused to an N-terminal fusion of mCherry and introduced them into mGBP2^−/-^ MEFs already expressing GFP-mGBP2. Cells were sorted using a BD FACSAriaIIu cell sorter and expression was confirmed via western blot using an anti-mCherry antibody. The subcellular localization of GFP-mGBP2 and the mCherry-tagged proteins of interest was assessed by confocal microscopy in IFN-γ-stimulated and unstimulated cells, under both uninfected and *T. gondii* infected conditions (Figs. 3, 4 and 5)

**Figure 3.**
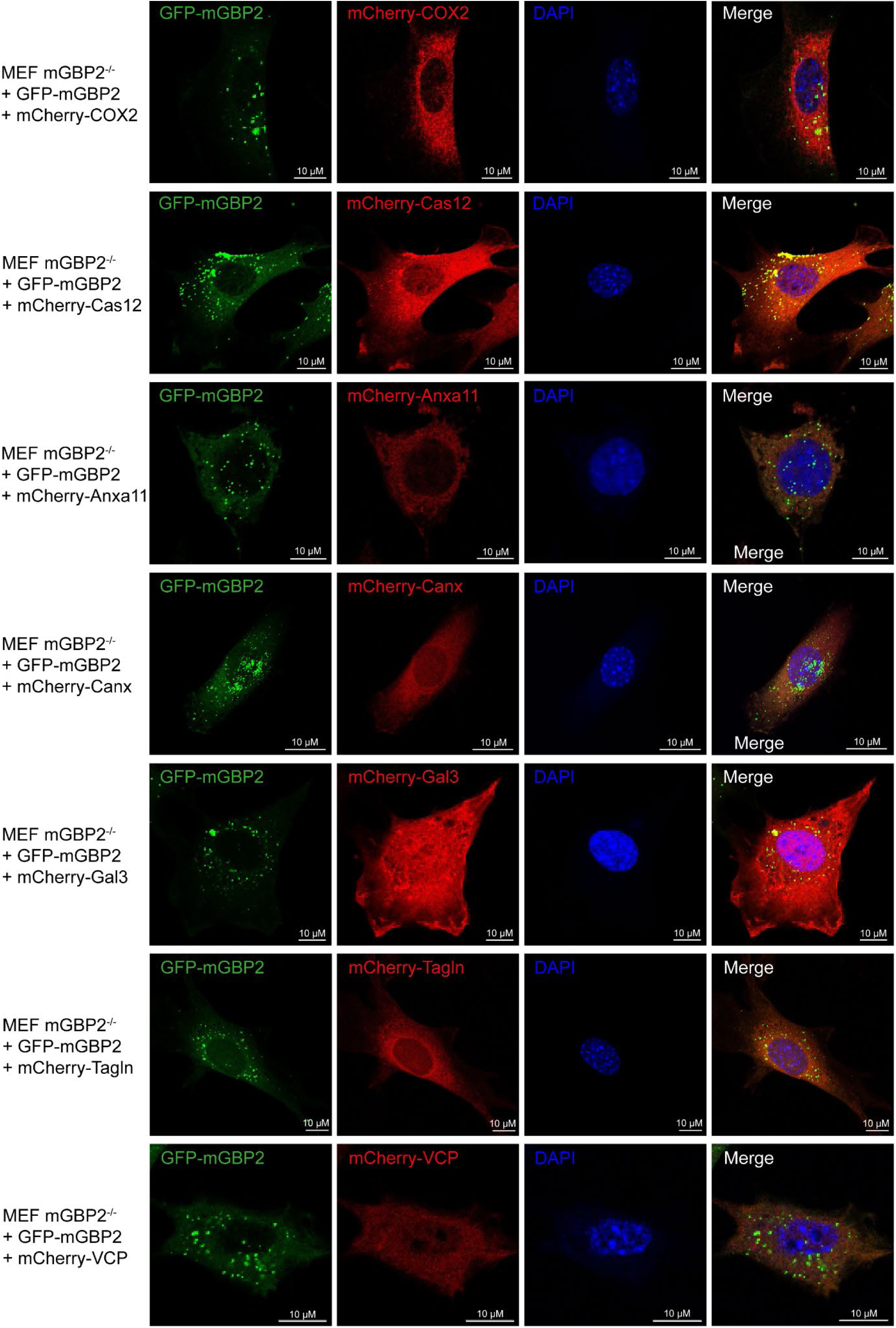
Intracellular localization of mGBP2 and potential interacting proteins. Localization was analyzed in GFP-mGBP2 and mCherry-fusion protein (COX2, Cas12, Anxa11, Canx, Gal3, Tagln, & VCP) co-transduced mGBP2-/- MEFs as indicated. Cells were treated with 200 U IFN-γ/ml for 16 h. After fixation, nuclei of cells were stained with DAPI. Slides were analyzed by confocal microscopy. Bars, 10 μm.

**Figure 4.**
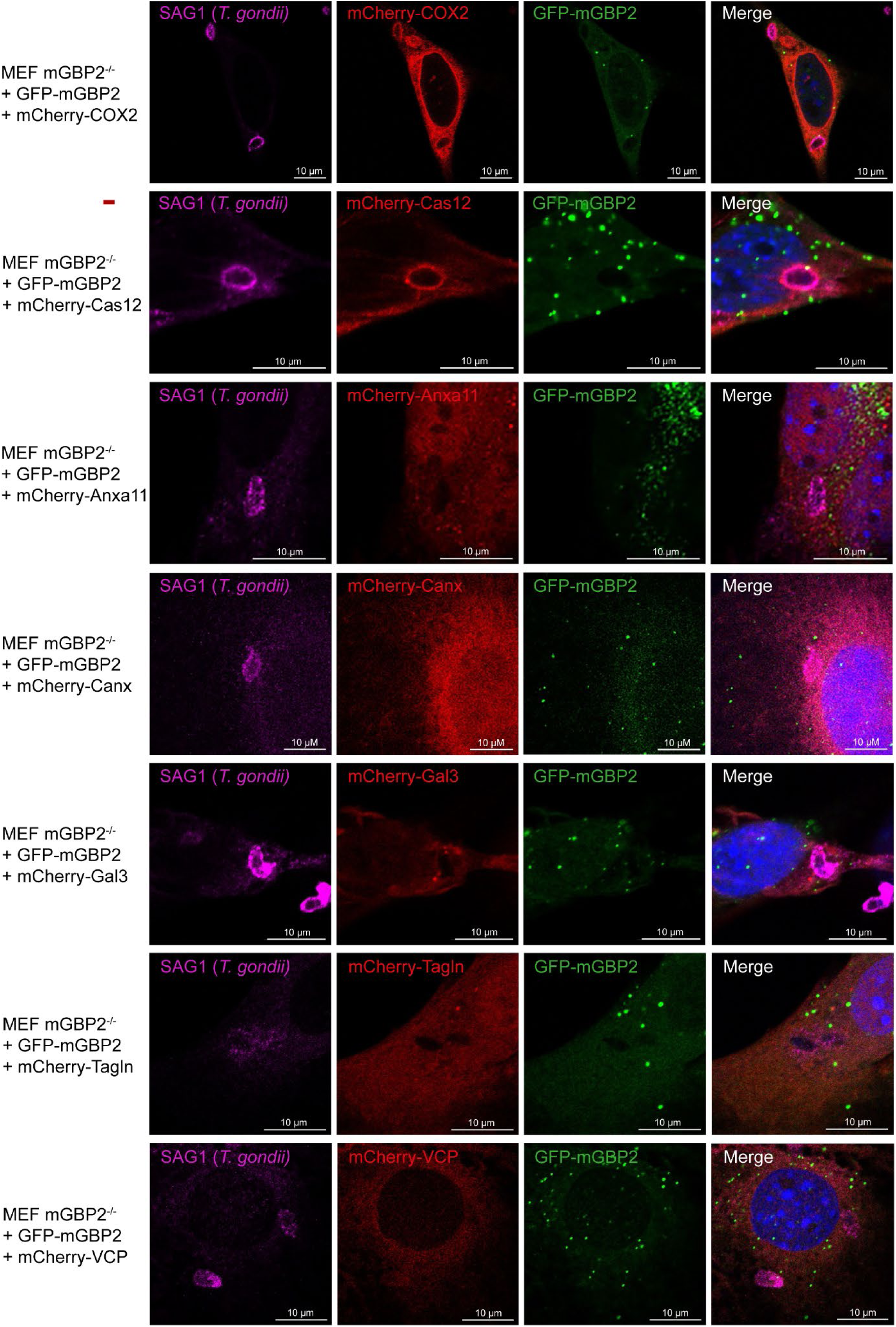
Intracellular colocalization of mGBP2 and potential interacting proteins at the PVM of *T. gondii.* Recruitment and colocalization was analyzed in GFP-mGBP2 and mCherry-fusion protein (COX2, Cas12, Anxa11, Canx, Gal3, Tagln, & VCP) co-transduced mGBP2^−/-^ MEFs as indicated. Cells infected with *T. gondii* for 2.5 h. After fixation, *T. gondii* was stained with an α-SAG1 antibody, and nuclei of cells were stained with DAPI. Slides were analyzed by confocal microscopy. Bars, 10 μm.

**Figure 5.**
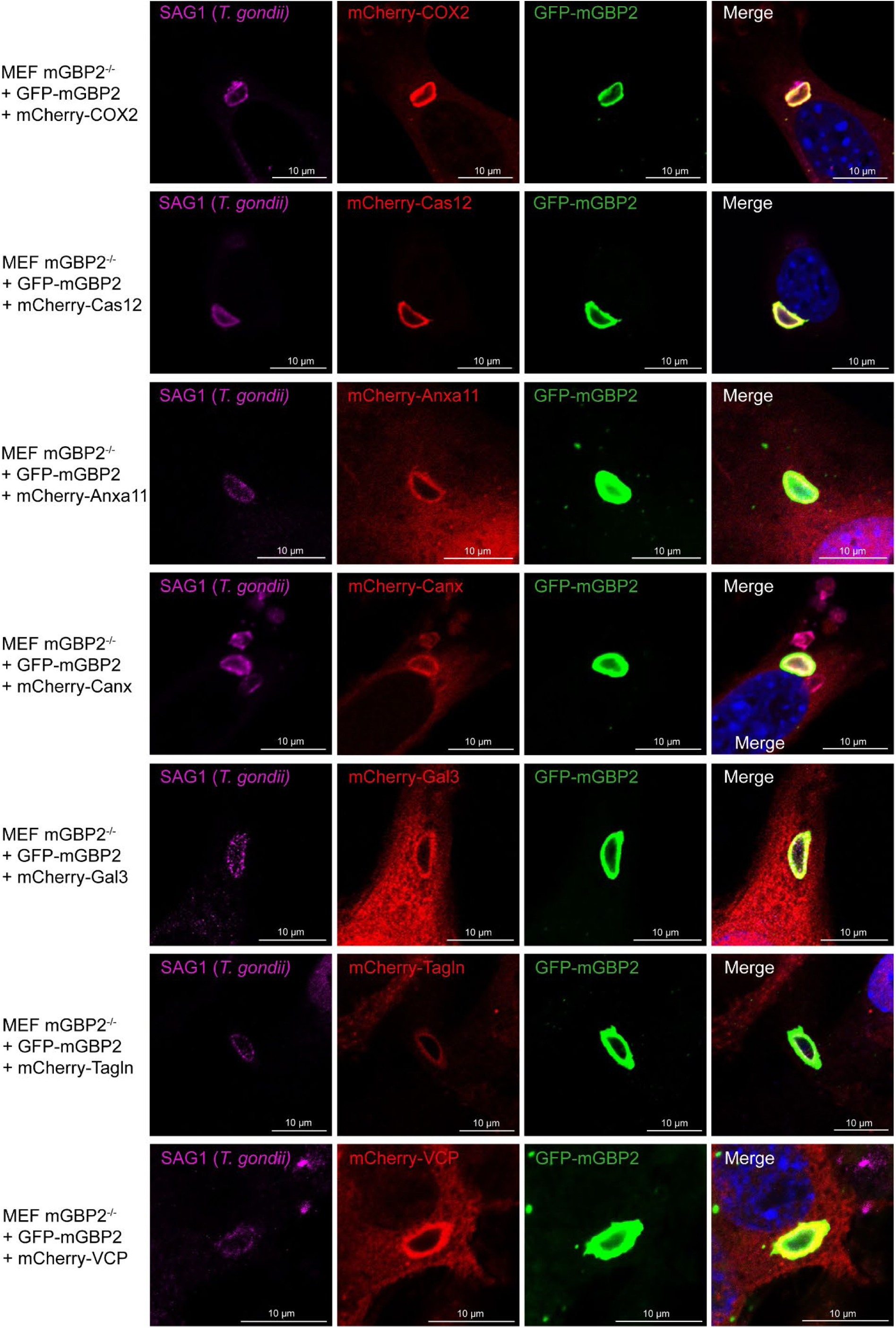
Intracellular colocalization of mGBP2 and potential interacting proteins at the PVM of *T. gondii*. Recruitment and colocalization was analyzed in GFP-mGBP2 and mCherry-fusion protein (COX2, Cas12, Anxa11, Canx, Gal3, Tagln & VCP) co-transduced mGBP2^−/-^ MEFs as indicated. Cells were treated with 200 U IFN-γ/ml for 16 h and subsequently infected with *T. gondii* for 2.5 h. After fixation, *T. gondii* was stained with an α-SAG1 antibody, and nuclei of cells were stained with DAPI. Slides were analyzed by confocal microscopy. Bars, 10 μm.

Firstly, we investigated the localization of the proteins of interest in IFN-γ stimulated but uninfected conditions (Fig. 3). COX2 showed typically localizations in the cytoplasm and ER that partially overlapped with mGBP2 localization, but no clear co-localization was detectable in uninfected cells. Cas12 could be detected in the cytosol and at the cytoplasmic side of the ER membrane, consistent with previous reports (Nakagawa, Zhu et al. 2000). Interestingly, this localization overlapped with mGBP2 in cytoplasmic VLS. Anxa11 was present in the cytosol showing no clear co-localization with mGBP2. Canx was described as an endoplasmic reticulum lectin (Myhill, Lynes et al. 2008). Consistently, it localized in the cytoplasm closely to the ER but did not show co-localization with mGBP2 in VLS. Gal3 could be found in both the cytoplasm as well as in the nucleus consistent with data previously described for human fibroblasts (Openo, Kadrofske et al. 2000). The localization of Gal3 partially overlapped with that of mGBP2. Tagln was localized in the cytosol as expected for an actin binding protein and could also be detected in the nucleus, however, Tagln did not co-localize with mGBP2 in uninfected conditions. VCP was localized in the cytosol showing no overlap with mGBP2 localization. However, a previously described localization of VCP in the nuclear compartment (Xing, Cai et al. 2025) could not be detected.

We then further analyzed the localization of the proteins of interest in *T. gondii* infection without IFN-γ stimulation. In these conditions mGBPs are not localized at the PV (Fig. 4). In the case of Anxa11, Canx, Gal3, Tagln and VCP no localization to *T. gondii* PVs could be seen in unstimulated conditions and without mGBP2 recruitment. From this, we conclude that either IFN-γ stimulation or the presence of mGBPs (including mGBP2) at the PV would be necessary for recruitment of these proteins. However, interestingly COX2 and Cas12 showed recruitment in unstimulated conditions indicating that recruitment of both proteins is independent of IFN-γ stimulation.

Finally, we investigated the localization and recruitment of the proteins of interest during *T. gondii* infection and IFN-γ stimulation (Fig. 5). In these conditions strong recruitment of mGBP2 to the PV can be detected as described previously (Degrandi, Kravets et al. 2013). After IFN-γ stimulation and *T. gondii* infection all investigated proteins i.e.; COX2/Ptgs2, Cas12, Anxa11, Canx), Gal3, Tagln, and VCP, showed recruitment to *T. gondii* PVs together with mGBP2. Apparently COX2 and Cas12 showed strongest recruitment compared to the other proteins as indicated by fluorescence intensities (not shown).

### COX2 and Cas12 are recruited to the *T. gondii* PV independently of mGBP2 expression and IFN-γ stimulation

For further analysis, we then investigated whether COX2 and Cas12 are able to recruit to *T. gondii* PVs in the absence of mGBP2 using mGBP2^−/-^ MEF cell lines expressing either mCherry-COX2 or mCherry-Cas12 (Fig. 6).

**Figure 6.**
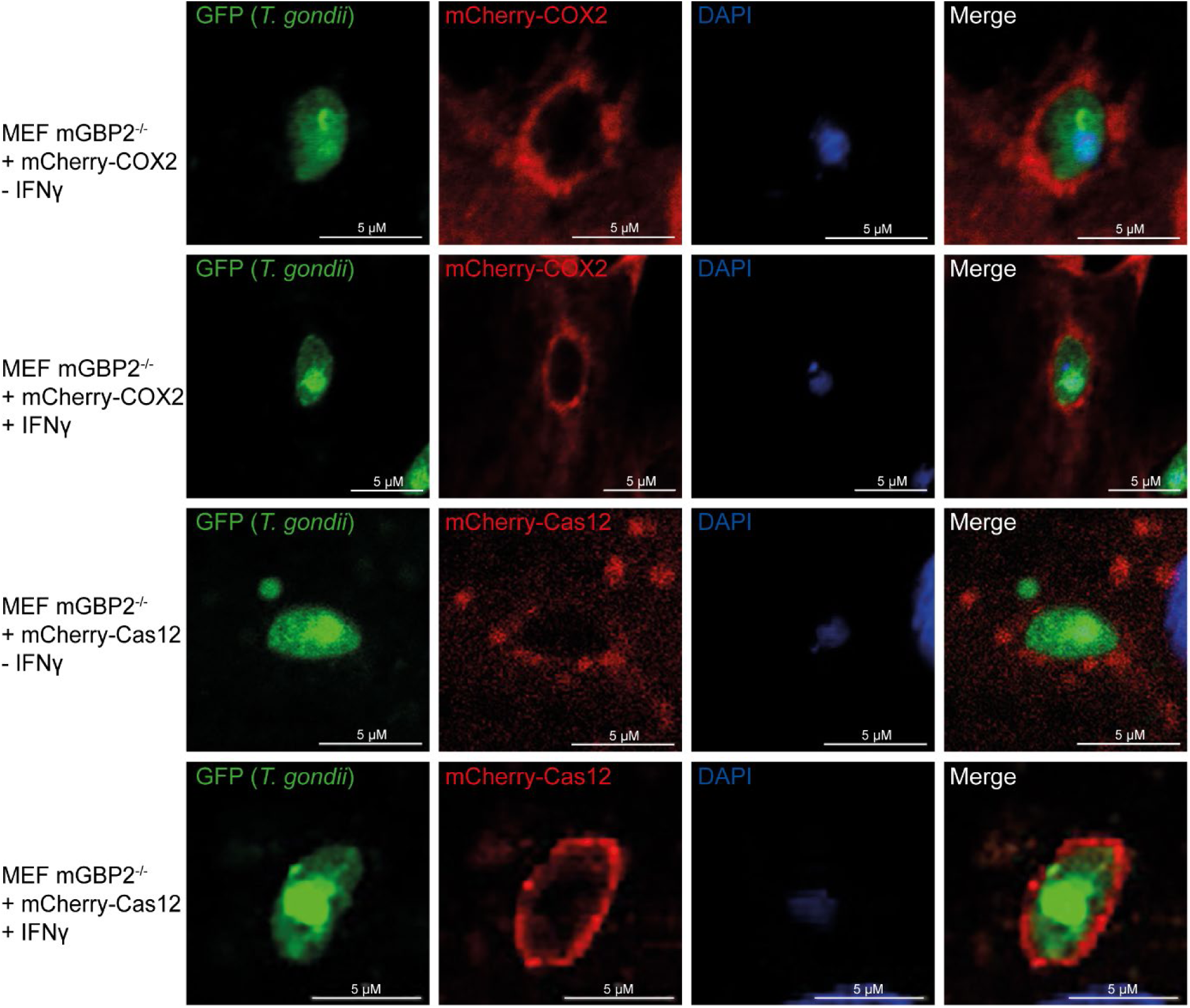
Intracellular localization of potential mGBP2 interacting proteins COX2 and Cas12 at the PVM of *T. gondii* in the absence of mGBP2. Recruitment and colocalization was analyzed in cherry-fusion protein (COX2, Cas12) transduced mGBP2^−/-^ MEFs. Cells were treated with IFN-γ for 16 h or left unstimulated and were subsequently infected with ME-GFP-Luc *T. gondii* for 2.5 h. After fixation, nuclei of cells were stained with DAPI. Slides were analyzed by confocal microscopy. Bars, 5 μm.

Analogous to the previous findings, both COX2 and Cas12 were able to recruit to *T. gondii* PVs independently of mGBP2 and without IFN-γ stimulation indicating a role of both proteins in *T. gondii* infection that is independent of mGBP2 recruitment.

### Expression of COX2, Cas12 and Gal3 is influenced by IFN-γ stimulation and *T. gondii* infection

The importance of IFN-γ signaling in *T. gondii* infection has been well described (Degrandi, Kravets et al. 2013, Yarovinsky 2014, Sasai and Yamamoto 2019). Particularly mGBPs like mGBP2 need IFN-γ stimulation to recruit to the *T. gondii* PV and lead to its disruption (Degrandi, Konermann et al. 2007, Yamamoto, Okuyama et al. 2012, Kravets, Degrandi et al. 2016). To investigate the role of IFN-γ on the expression of each of the proteins of interest we used RT-PCR to determine their regulation after IFN-γ or LPS stimulation and *T. gondii* infection (Fig. 7).

**Figure 7.**
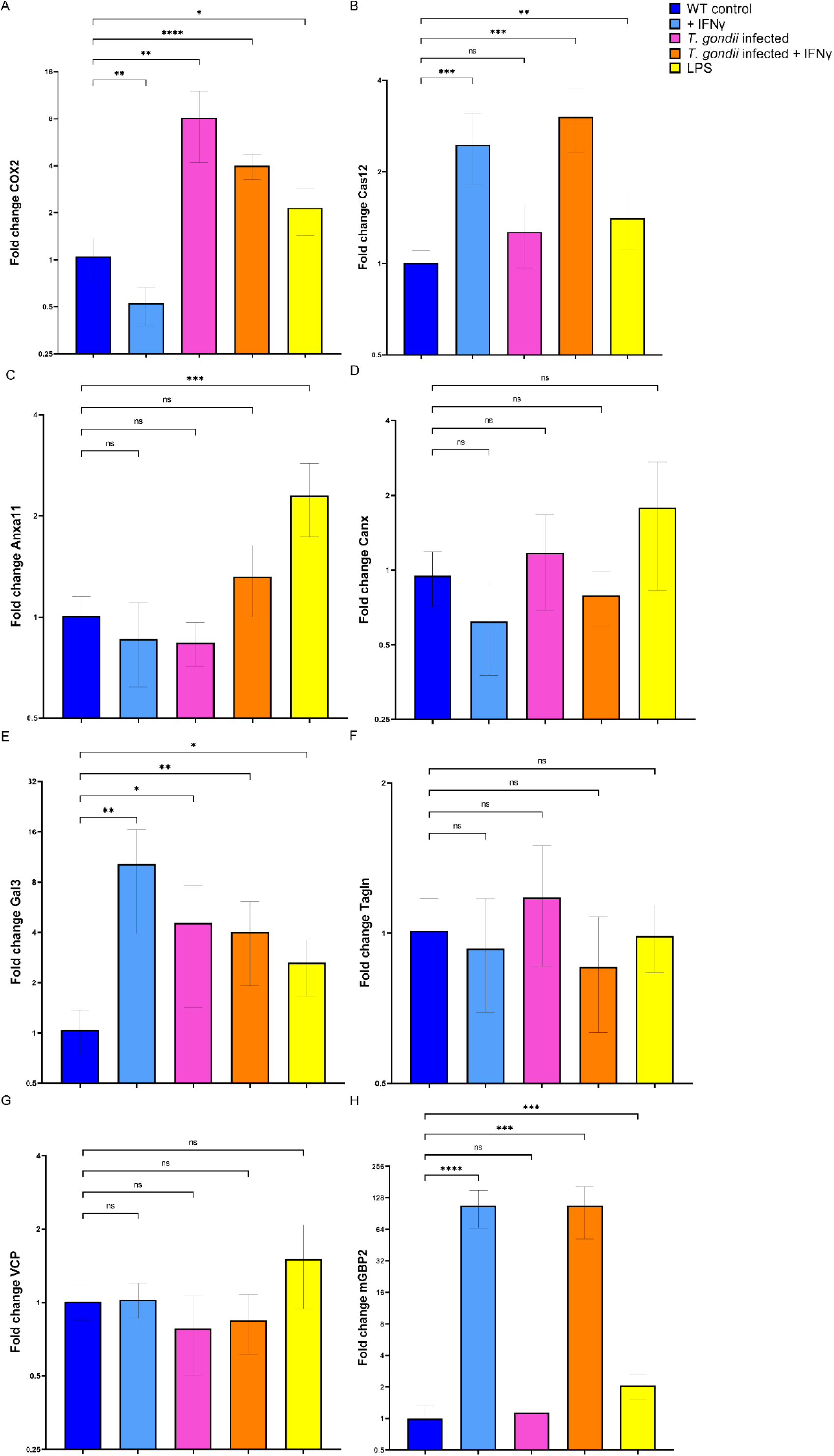
Fold change of mRNA expression of proteins of interest after IFN-γ or LPS stimulation and *T. gondii* infection. WT MEF cells were either stimulated with 200 U/ml IFN-γ or 500 ng/ml LPS for 16 h and left uninfected or subsequently infected with *T. gondii* for 2.5 h. RNA was isolated from the cells and transcribed into cDNA followed by Real-time. mGBP2 was used as a control for IFN-γ induction. Fold change was normalized using beta-actin. Data shown are from the means of three independent experiments each performed in triplicates (n = 3) ± SD. One-Way ANOVA was performed with Geisser-Greenhouse correction and Dunnet-Test with each sample compared to mean value of wildtype control.

Interestingly, qRT-PCR showed that the expression of COX2 was reduced after IFN-γ stimulation (Fig.7A), which was confirmed by Western Blot analysis (Fig. 8). On the other hand, COX2 expression was significantly increased after LPS stimulation, as has been described previously (Giuliano and Warner 2002). In *T. gondii* infection, the expression of COX2 was drastically increased confirming COX2 as an inflammatory and inducible isoform of cyclooxygenases. The decrease of COX2 expression after IFN-γ stimulation was, on the first view, somewhat surprising since in the literature an opposite effect in breast cancer cells has been described (Cheng, Ridnour et al. 2023)). However, other studies showed that the effect of IFN-γ on COX2 can be highly species and cell type dependent (Minghetti, Polazzi et al. 1996, Deng, Montero et al. 2007).

**Figure 8.**
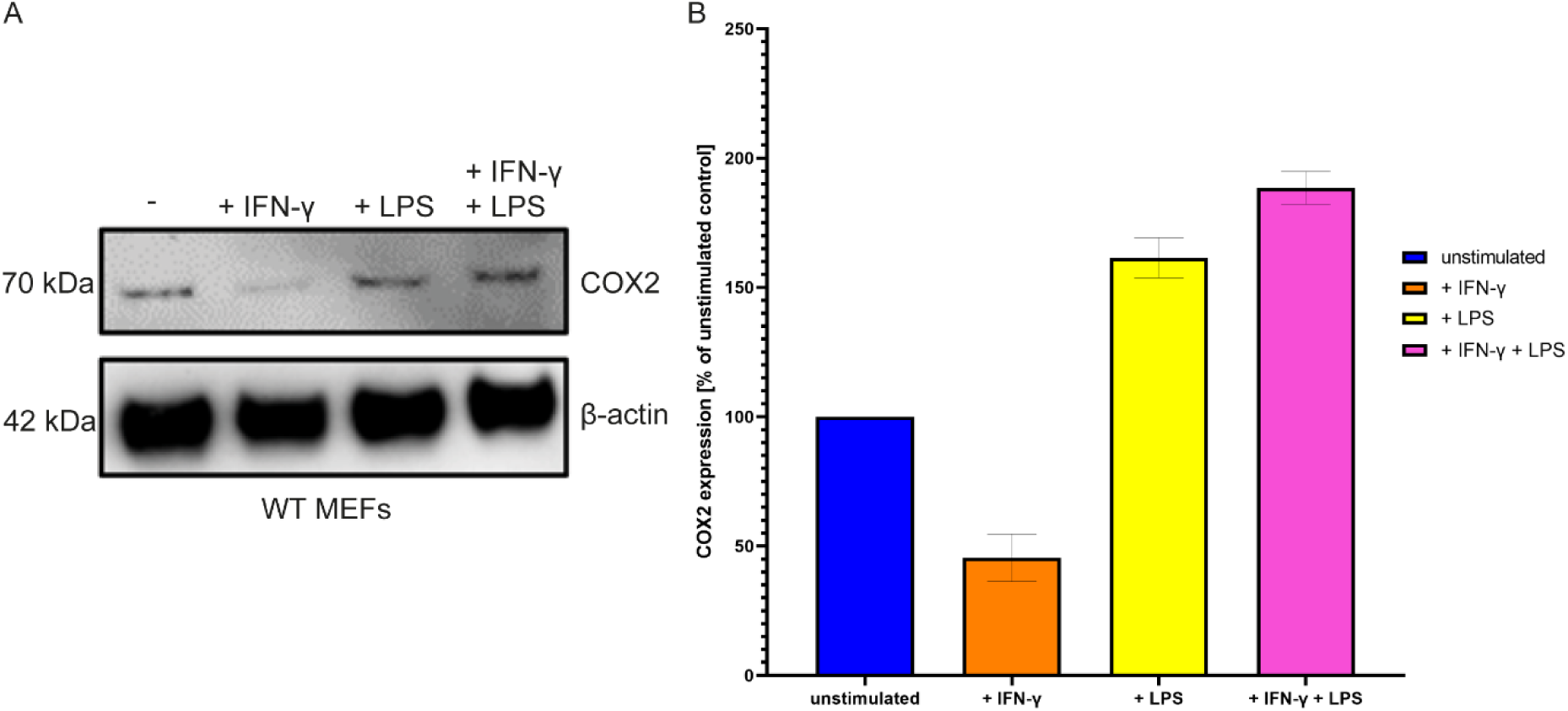
2 – (A) COX2 expression in wildtype MEF cells as determined by western blot. Cells were stimulated with 200 U IFN-γ or 500 ng/ml LPS or both for 16 h. COX2 was detected using a COX2 antibody (upper panel). As loading control an anti β-actin antibody was used (lower panel). (B) COX2 expression amounts were quantified using ImageJ and normalized to the β-actin control. COX2 expression changes relative to the unstimulated control cells means of two independent experiments are depicted.

Cas12 has previously been described as an IFN-γ inducible caspase (Kalai, Lamkanfi et al. 2003) which we could be confirmed using qRT-PCR (Fig. 7B). However, *T. gondii* infection did not have an effect on Cas12 expression. Interestingly, Cas12 was also increased upon LPS stimulation. Anxa11 showed no changes in expression upon IFN-γ stimulation or *T. gondii* infection. A slight increase in expression could be observed after LPS stimulation (Fig. 7C). Canx showed no change in expression in all four stimulation conditions which was expected due to it serving as a house keeping gene and being essential for ER function and correct glycoprotein folding (Schrag, Bergeron et al. 2001) (Fig. 7D). Gal3 showed increased expression after all three stimulus conditions (Fig. 7E). VCP and Tagln also showed no expression change in all conditions (Fig. 7G). mGBP2 expression was used as control for IFN-γ stimulation showing a drastic increase of expression as has been previously described (Fig. 7H) (Boehm, Guethlein et al. 1998, Degrandi, Konermann et al. 2007).

### Inhibition of COX2 reduces *T. gondii* proliferation

To investigate the function of COX2 as an effector in *T. gondii* infection we inhibited its function using the COX2 enzyme inhibitor Celecoxib and performed *T. gondii* proliferation assays in MEF and HS27 cells. Celecoxib (Celebrex®; Pfizer Inc.) is a selective COX2 inhibitor that was released onto the market in 2000 for the symptomatic treatment of arthritis. Celecoxib inhibits the conversion of arachidonic acid to prostaglandin H2 by binding reversibly to the active site of COX2 (Clemett and Goa 2000)..

Interestingly, in both human as well as in murine fibroblasts the inhibition of COX2 function led to reduced *T. gondii* proliferation with IC_50_ values in the low micromolar concentrations (5.34 ± 0.12 µM and 8.82 ± 2.93 µM, respectively) while the cytotoxic effect on the host cells was only present in higher concentrations above 50 µM (CC_50_ = 132.9 µM and 89.1 µM, respectively) (Fig. 9). This indicates a positive effect of COX2 function in the parasite’s survival and growth. A similar effect has been described in trophoblasts and villious explants explants (de Souza, Silva et al. 2021). The non-approved COX2 inhibitor Etoricoxib showed a similar effect in human fibroblasts (results not shown).

**Figure 9.**
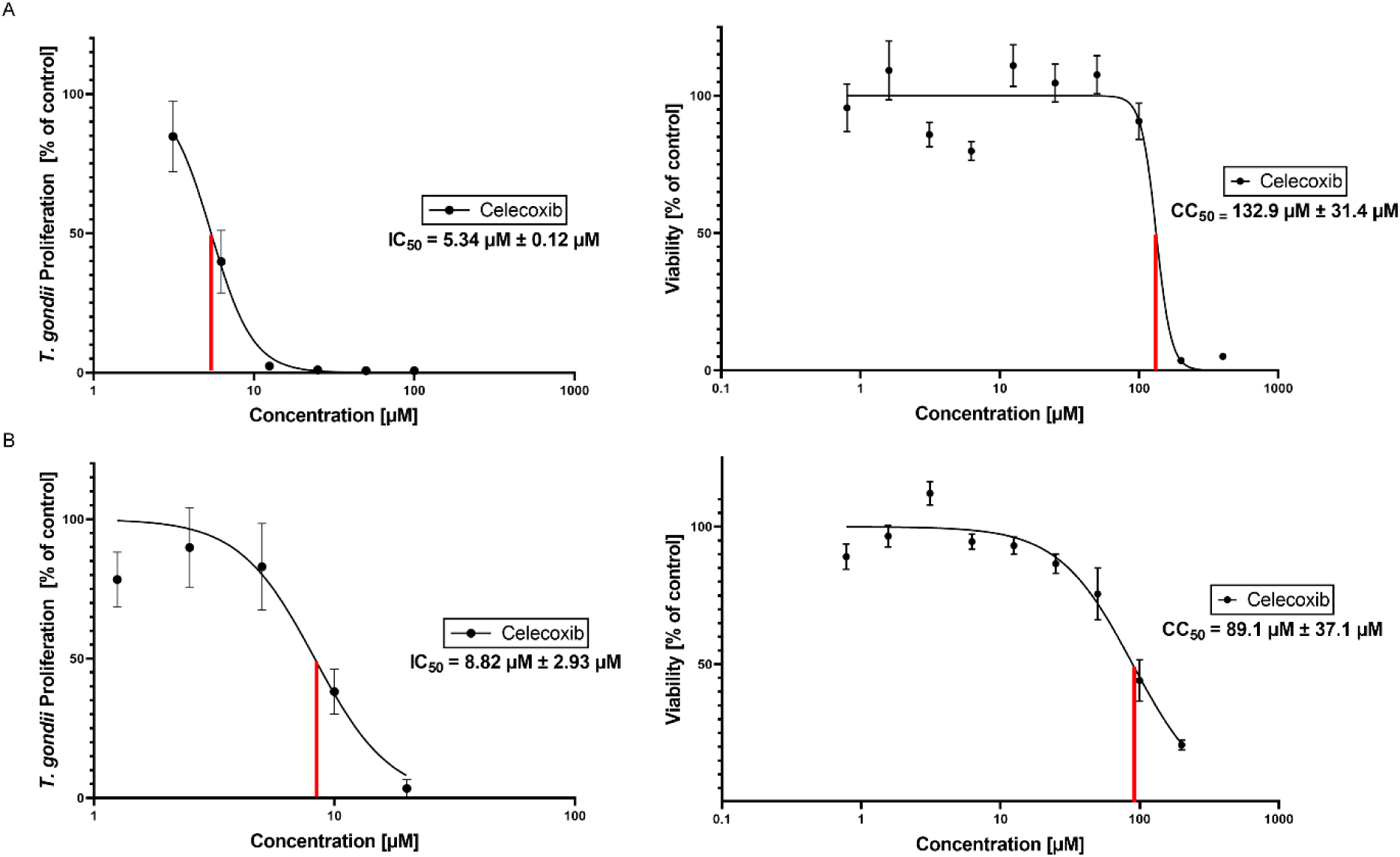
Anti-toxoplasma activity and cytotoxicity of the COX2 inhibitor Celecoxib on human fibroblasts Hs27 (A) and MEF cells (B). The inhibitory activities were determined by *T. gondii* proliferation *in vitro* inhibition using ME-GFP-Luc *T. gondii*. Cytotoxicity was measured by MTT assays on human fibroblasts Hs27 or MEF cells. Data shown are from the means of three independent experiments, each performed in triplicates (n = 3) ± SEM.

### The cyclooxygenase isoform COX1 does not recruit to *T. gondii* PVs

As COX2 is the inducible isoform of cyclooxygenase, we aimed to also investigate the endogenously expressed COX1 isoform (Seibert, Zhang et al. 1997). In contrast to COX2, COX1 was not enriched in *T. gondii*-infected cells compared with uninfected cells in MS analysis following APEX2 proximity labeling (Fig.2, Suppl. Table 1). Notably, in contrast to COX2, we could not detect COX1 recruitment to *T. gondii* PVs in the presence or absence of IFN-γ stimulation (Fig. 10A). Subsequently, we tested the effect of the selective COX1 inhibitor SC560 (COX-1 IC_50_ = 0.009 μM; COX-2 IC_50_ = 6.3 μM) (Smith, Zhang et al. 1998) on *T. gondii* proliferation (Fig. 10).

**Figure 10.**
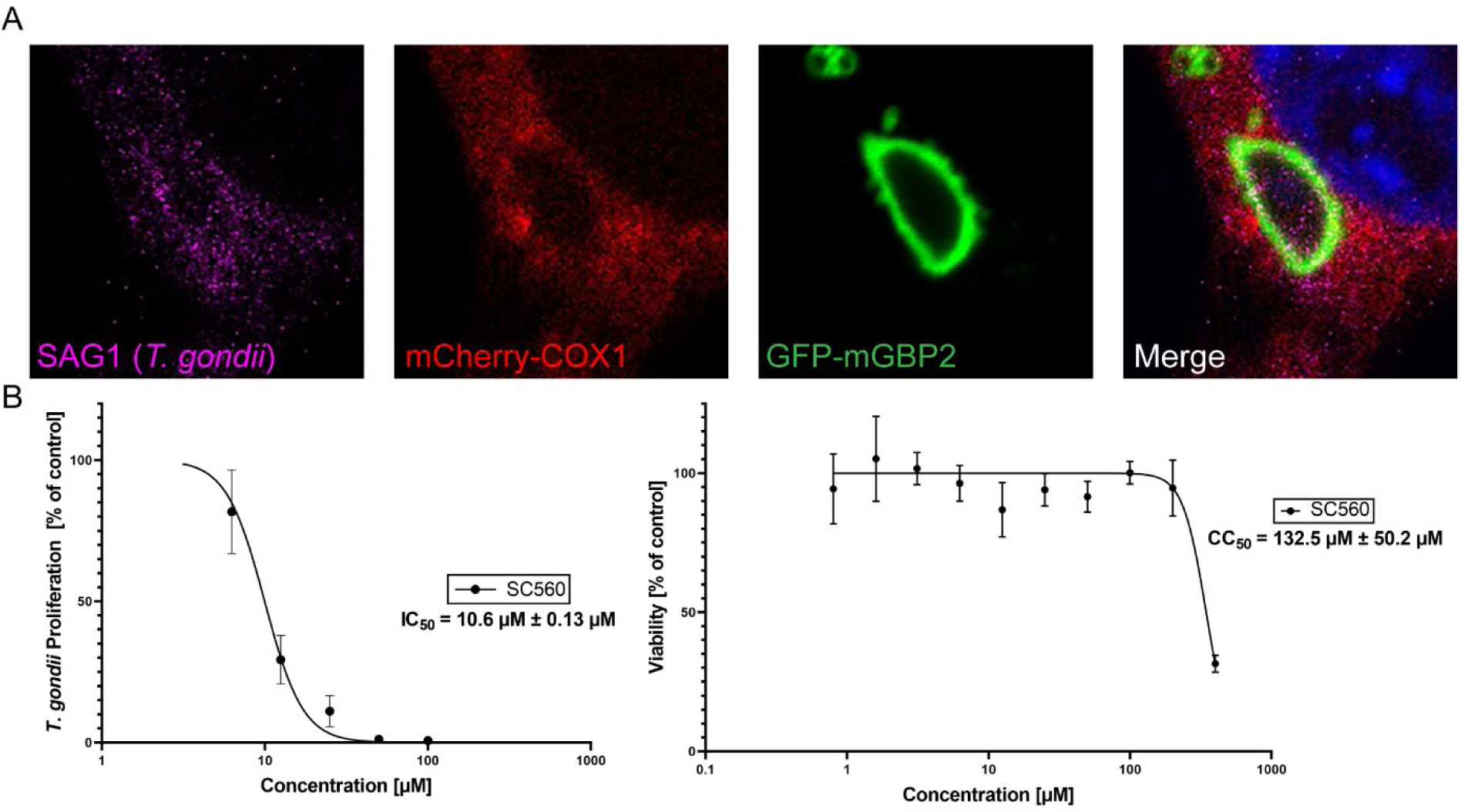
Investigation of the cyclooxygenase isoform COX1. (A) Intracellular localization of mGBP2 and Cox1 at the PV of *T. gondii.* Recruitment and colocalization was analyzed in GFP-mGBP2 and mCherry-Cox1 expressing mGBP2^−/-^ MEFs. Cells were stimulated with IFN-γ for 16 h and subsequently infected with *T. gondii* for 2.5 h. After fixation, *T. gondii* was stained with an α-SAG1 antibody, and nuclei of cells were stained with DAPI. Slides were analyzed by confocal microscopy. Bars, 10 μm. (B) Anti-toxoplasma activity and cytotoxicity on human fibroblasts Hs27 of the COX1 inhibitor SC560. The inhibitory activities were determined by inhibition of the *T. gondii* proliferation *in vitro* using ME-GFP-Luc *T. gondii*. Cytotoxicity was measured by MTT assays on human fibroblasts Hs27. Data shown are from the means of three independent experiments each performed in triplicates (n = 3) ± SEM.

Although the COX1 inhibitor SC560 showed reduction in *T. gondii* proliferation, the IC_50_ (10.6 µM) falls within the range at which SC560 also inhibits COX2. Therefore, the *T. gondii* inhibitory effect cannot be attributed specifically to COX1. SC560 showed no cytotoxic effects in the range of inhibitory concentrations (Fig. 10B). We also investigated the effect of IFN-γ stimulation*, T. gondii* infection, and LPS stimulation on the expression levels of COX1.

In contrast to COX2 the expression of COX1 did not change significantly upon all conditions (Fig. 11).

**Figure 11.**
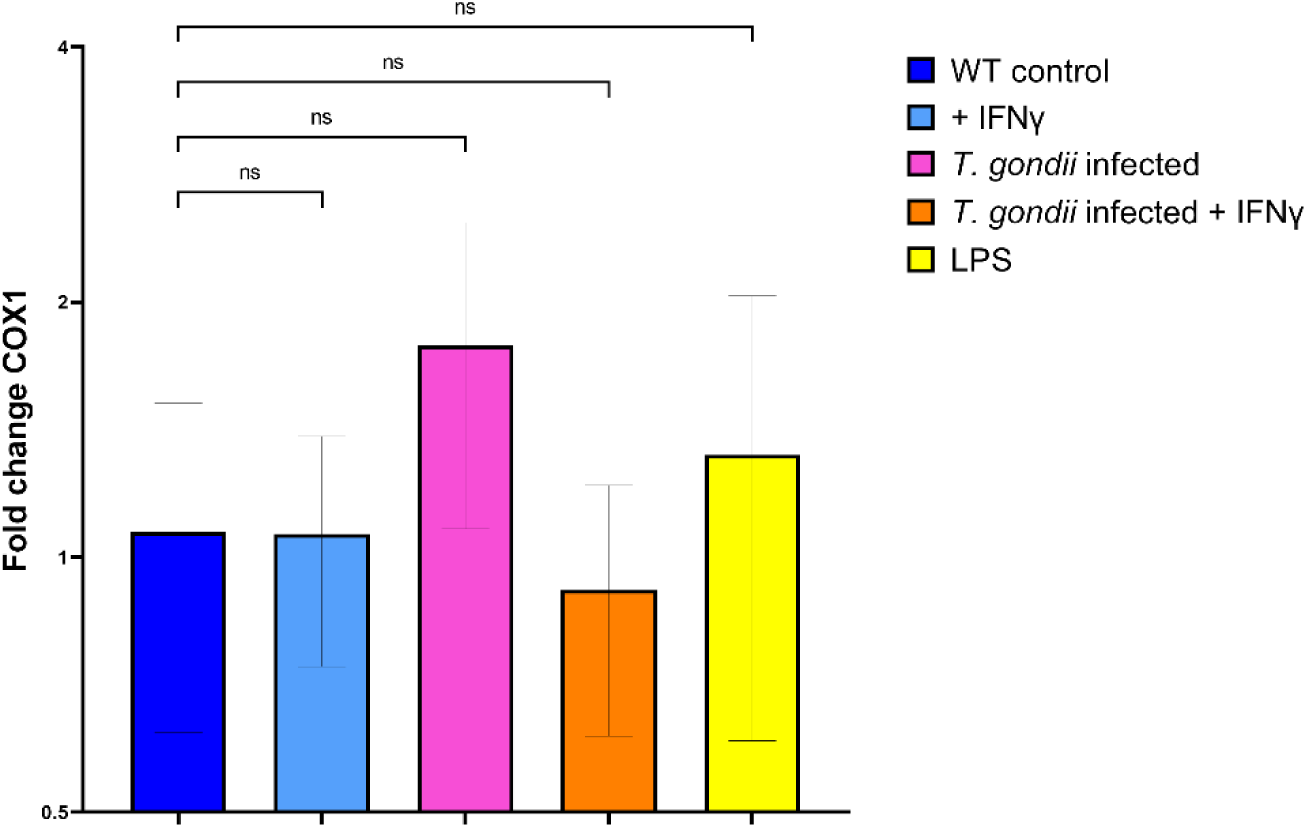
Fold change of COX1 mRNA after IFN-γ or LPS stimulation and *T. gondii* infection. WT MEF cells were either stimulated with 200 U/ml IFN-γ or 500 ng/ml LPS for 16 h and subsequently infected with *T. gondii* for 2.5 h. RNA was isolated from the cells and transcribed into cDNA followed by Real-time PCR. Fold change was normalized using beta-actin. Data shown are from the means of three independent experiments each performed in triplicates (n = 3) ± SD.

In conclusion, seven proteins (COX2, Cas12, Anxa11, Canx, Gal3, Tagln, VCP) that localize to PVs together with mGBP2 and display distinct functional roles during infection were identified. COX2 and Cas12 appear to act independently of IFN-γ stimulation and mGBP2 recruitment, with COX2 promoting *T. gondii* survival, which could be confirmed by use of COX2 inhibitors. Expression analysis further revealed that COX2 is upregulated during *T. gondii* infection but downregulated during IFN-γ stimulation, whereas Cas12 and Gal3 expression are induced by IFN-γ. Anxa11, Canx, Tagln and VCP showed no changes in expression after IFN-γ stimulation. Collectively, this study used an APEX2-based proximity labelling approach to identify an important set of proteins important for the cell autonomous immunity and for intracellular survival of *T. gondii* parasites that can be exploited for treatment of *T. gondii* infections.

## DISCUSSION

By mapping the protein environment of mGBP2 at the *T. gondii* PV, our APEX2-based proximity labeling study expands the repertoire of candidate PV-associated host factors and provides a basis for a deeper understanding of the cellular processes involved in IFN-γ-induced cell-autonomous immunity.

Two of these proteins, Cas12 and COX2, were found to recruit to the *T. gondii* PV independently of IFN-γ stimulation or the presence of mGBP2. However, IFN-γ drastically increased the expression of Cas12 while *T. gondii* infection alone did not lead to changes in Cas12 transcripts. Cas12 is phylogenetically related to the inflammatory caspase family, which comprises caspase 1 (Cas1), 4, and 5 in humans and caspase 1 and 11 in mice. Cas1, the central inflammatory caspase, is activated within the inflammasome, a large complex formed by Nod-like receptors (NLRs), in response to pathogen invasion or host-derived danger signals (Petrilli, Dostert et al. 2007). The activity of Cas1 is regulated by related inflammatory caspases, including Cas12 which represses inhibits NF-κB pathways and the inflammasome via Cas1 catalysis (Saleh, Mathison et al. 2006, Roy, Sharom et al. 2008). In previous studies murine Cas12 was shown to limit the immune control of *Plasmodium* replication whereas Cas12 deficiency leads to hyperactivation of NF-κB and enhanced IFN-γ production (Saleh, Mathison et al. 2006, Labbe, Miu et al. 2010). The knowledge on the role of Cas12 in *T. gondii* infection is limited up to now. *In vitro* and *in vivo* studies suggested an involvement of Cas12 in apoptosis regulation in neuronal tissue and cells upon *T. gondii* infection (Wang, Zhou et al. 2014, Wan, Gong et al. 2015). However, based on our results, an immunoregulatory function of Cas12 in *T. gondii* infection appears feasible, since we observed early recruitment to PVs without IFN-γ dependent activation of antimicrobial cell autonomous immunity. Further studies are needed to confirm these hypotheses.

Recently, we showed that Gal9 colocalizes to the *T. gondii* PV together with mGBP2 and that. Gal9 contributes to mGBP2-dependent restriction of *T. gondii* growth (Kravets, Poschmann et al. 2025). Interestingly, in this study we could demonstrate that also Gal3 is present at the PV after IFN-γ stimulation together with mGBP2. Moreover, IFN-γ stimulation as well as *T. gondii* infection increased the expression of Gal3 in MEF cells. Galectins represent a subgroup of lectins that contain one or two carbohydrate-recognition domains (CRDs), which exhibit an affinity for glycans containing β-galactoside (Liu and Rabinovich 2010). Gal3 participates in several biological processes. In particular Gal3 plays a key role in immune responses, the regulation of immune tolerance and also in responses associated with host-pathogen interactions (Sato, St-Pierre et al. 2009, Liu and Rabinovich 2010). Importantly, Gal3 has been implicated in the innate immune response to *T. gondii* infection, where Gal3-deficient mice showed reduced inflammatory infiltration in several organs but increased parasite burden in the brain. This was accompanied by enhanced IL-12 production by dendritic cells and a stronger Th1-polarized response, suggesting that Gal3 contributes both to inflammatory cell recruitment and to the regulation of adaptive immunity (Bernardes, Silva et al. 2006). Furthermore, extracellular Galectins can bind directly to bacteria and mediate bacteriostatic or bactericidal effects (Chen, Weng et al. 2014). In this context, Gal3 has been described as a sensor of vacuolar rupture that ensues upon invasion of bacteria into the host cell (Paz, Sachse et al. 2010). Interestingly, previously Gal3 was shown to direct mGBP2 and mGBP1 to intracellular vacuoles containing *Y. pseudotuberculosis* and *L. pneumophila* (Feeley, Pilla-Moffett et al. 2017). Whether Gal3 may adopt a similar function in *T. gondii* infection has yet to be explored.

VCP/p97 is a hexameric AAA+ ATPase involved in membrane fusion and ubiquitin-dependent protein processing (Ye 2006). VCP and its cofactor UBXD1 recruit to *T. gondii* PVs in IFN-γ-stimulated human endothelial cells. VCP recruitment also occurs in other human cell types (Clough, Fisch et al. 2023). In our study, we extend these observations to murine cells, where VCP colocalized with mGBP2 after IFN-γ stimulation without significant changes in *Vcp* transcripts. Whether VCP processes ubiquitinated proteins at the PV remains unresolved.

Anxa11 is a calcium-dependent phospholipid-binding protein that couples protein and lipid phase transitions on lysosomal membranes (Nixon-Abell, Ruggeri et al. 2025). It interacts with ALG-2/PDCD6 (Satoh, Shibata et al. 2002), which links it to Sec31A (Shibata, Kanadome et al. 2015) and binds ALIX/PDCD6IP (Missotten, Nichols et al. 1999), an ESCRT-associated regulator (Khan, Alaamery et al. 2020). Sec31A and ALIX were enriched in our samples; ALIX was previously localized to PVs in human cells (Cygan, Jean Beltran et al. 2021). Although Anxa11’s role at the *T. gondii* PV remains unresolved, Anxa1 is upregulated in ocular toxoplasmosis (Mimura, Tedesco et al. 2012), and an Anxa1-derived peptide reduced parasitism in human placental explants (de Oliveira Cardoso, Moreli et al. 2018).

Calnexin (Canx) is an ER membrane lectin chaperone involved in glycoprotein folding and quality control (Paskevicius, Farraj et al. 2023). Its PV-associated localization may reflect apposition of host ER, which *T. gondii* recruits alongside other organelles (Coppens and Romano 2018). This association requires further spatial characterization.

Tagln (transgelin, also known as smooth muscle protein 22α, SM22α) is an actin-binding protein of the calponin family (Assinder, Stanton et al. 2009). Its PV-associated localization could relate to cytoskeletal remodeling, as described during *T. gondii* infection of dendritic cells (Weidner, Kanatani et al. 2013), although this connection remains to be tested.

Among all mGBP2-proximal protein candidates identified in this study, COX2 showed the highest abundance in *T. gondii* infected cells. COX2 is an enzyme responsible for catalyzing the production of prostaglandins (PGEs) from arachidonic acid at the site of inflammation. Arachidonic acid is found within the cell membrane and is further transformed by Cyclooxygenases into eicosanoids (Giuliano and Warner 2002, Kalinski 2012). Two isoforms of COX have been identified. COX1 is constitutively expressed in most cell types while COX2 is induced by inflammatory mediators, mainly cytokines, and remains mostly undetectable in most unstimulated mammalian cells (Giuliano and Warner 2002). PGE_2_ exerts immunosuppressive and vasodilatory activities and enhances vascular permeability, thus facilitating pathogen invasion. It has been shown that various infection by pathogens, including *T. gondii*, of human cells lead to an upregulation of mRNA and protein levels for COX2 (Kim, Kim et al. 2000, Kim, Ahn et al. 2006, Moraes, Diniz et al. 2015, Martinez-Colon and Moore 2018). It has been observed that *T. gondii* infection leads to an increase of lipid droplets in mice peritoneal macrophages (Mota, Roberto Neto et al. 2014) as well as COX2 and PGE_2_ expression in cultures of skeletal muscle cells (Gomes, Magalhaes et al. 2014). Furthermore, it has been shown that COX2 inhibition upregulates pro-inflammatory cytokines, reduces lipid droplet availability and decreases intracellular proliferation of *T. gondii* (Pereira, Silva et al. 2019, de Souza, Silva et al. 2021). Our results showed that COX2 is present at the *T. gond*ii PV independently of IFN-γ stimulation or the presence of mGBP2. Using inhibitors, we showed that COX2 function is important for *T. gondii* growth and proliferation in human and murine fibroblasts, indicating a beneficiary effect of COX2 on *T. gondii* survival, most likely by providing the parasite with necessary lipids and other metabolites. This supportive role for parasite survival was further confirmed by a high induction of COX2 expression upon *T. gondii* infection. Meanwhile IFN-γ stimulation led to a decreased expression indicating an antagonistic play between IFN-γ induced response against infection by the host cell and *T. gondii* induced protection against the innate immune response. We could also confirm that these effects are specific to COX2 and do not extend to the COX1 isoform.

Concluding, we identified seven proteins of interest that localize to *T. gondii* PVs together with mGBP2. These proteins of interest showed different properties with COX2 and Cas12 acting independently of IFN-γ stimulation and mGBP2 recruitment and most likely providing a positive effect on *T. gondii* survival. Meanwhile Gal3 and VCP could display a protective role against infection of the host cell. The presence of Anxa11, Tagln and Canx is likely explained by changes in the cell environment upon *T. gondii* infection such as rearranging the cytoskeleton and recruitment of ER to provide the parasite with necessary metabolites. Overall, all proteins of interest display interesting functions for further analysis to understand the molecular environment present at *T. gondii* PV.

**Table 1.**
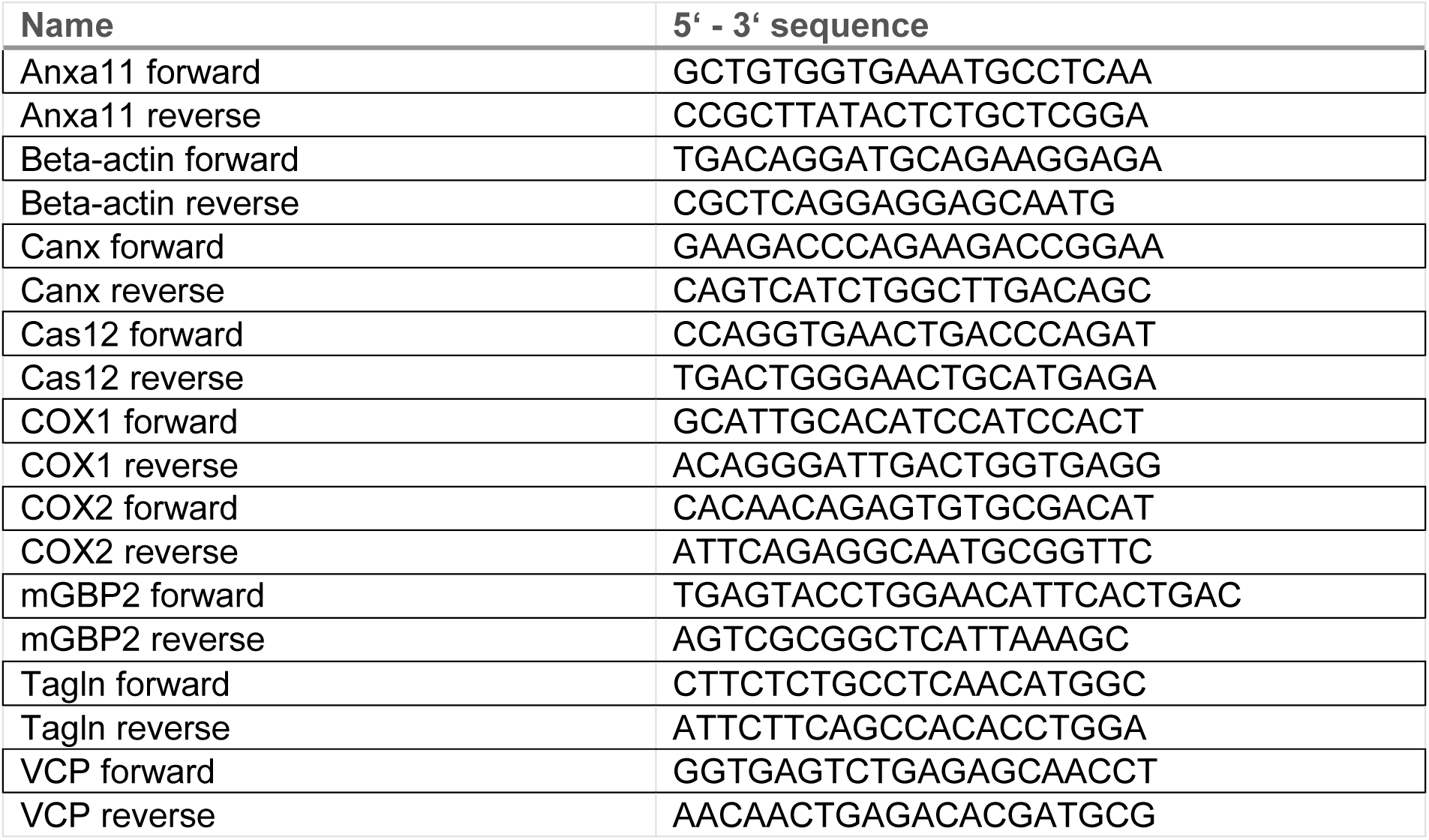
Primers used for qPCR.

## ACKNOWLEDGEMENTS

We gratefully acknowledge the generous support of the DFG (GRK2158, SFB 1208 Project-ID 267205415) and Jürgen Manchot Foundation We thank Karin Buchholz, Nicole Küpper and Julia Mock for their technical assistance.

